# Reactive Oxygen species dependent increase in H3K27 acetylation by intermittent hypoxia is regulated by H3S28 phosphorylation

**DOI:** 10.1101/2024.09.09.612097

**Authors:** Ning Wang, Matthew Hildreth, Nanduri R. Prabhakar, Jayasri Nanduri

**Affiliations:** Institute for Integrative Physiology and Center for Systems Biology of O2 Sensing, The University of Chicago, Chicago, IL

**Keywords:** Histone acetylation, phosphorylation, intermittent hypoxia, reactive oxygen species, histone deacetylases histone acetylases

## Abstract

Histones play a crucial role in regulating gene expression through post –translational modifications (PTMS) which include acetylation, methylation and phosphorylation. We have previously identified histone 3 acetylation (H3Kac) and methylation (H3Kme) as an early epigenetic mechanism associated with intermittent hypoxia (IH), a hallmark feature of sleep apnea. The goal of the present study was to determine the molecular mechanisms underlying IH increased H3 acetylation. IH-induced H3 acetylation was blocked by an antioxidant. Conversely, reactive oxygen species (ROS) mimetics, increased H3 acetylated protein expression similar to IH, suggesting a role for ROS. Trichostatin A (TSA), an HDAC (histone deacetylase) inhibitor mimicked IH-induced H3 acetylation under normoxic conditions, while pharmacological blockade of p300/CBP (HAT, histone acetylase) with CTK7A abolished IH-induced H3 acetylation. These results suggest that interplay between HATs and HDACs regulate ROS-dependent H3 acetylation by IH. Lysine 27 (H3K27) on H3 was one of the lysines specifically acetylated by IH and this acetylation was associated with dephsophorylation of H3 at serine 28 (H3S28). Inhibition of S28 dephosphorylation by protein phosphatase inhibitors (PIC or Calyculin A), prevented H3K27 acetylation by IH. Conversely, inhibiting K27 acetylation with CTK7A, increased S28 phosphorylation in IH-exposed cells. These findings highlight the intricate balance between H3 acetylation and phosphorylation in response to IH, shedding light on epigenetic mechanism regulating gene expression. (Supported by NIH-PO1-HL90554).

## INTRODUCTION

Obstructive sleep apnea patients experience recurrent apneas, characterized by transient, repetitive cessations of breathing. OSA patients exhibit elevated sympathetic excitation and blood pressures (1–3). Intermittent hypoxia (IH) resulting from apneas is the primary stimulus for evoking sympathetic excitation (2, 4, 5). Hypoxia inducible factor (HIF-1)–dependent transcriptional activation of the pro-oxidant NADPH oxidase *(NOX)* genes, and the subsequent ROS signaling is a major cellular mechanism underlying IH-induced sympathetic nerve activation (6–8).

Epigenetic mechanisms involving post-translational modifications (PTM) of the amino terminal tails of histone proteins play a crucial role in gene regulation. Specifically, histone acetylation acts as an epigenetic marker shaping chromatin and influencing gene activity (9, 10) Histone hyperacetylation is associated with induction of gene expression while hypoacetylation, is associated with gene repression (11, 12). The acetylation status of histones, dictated by a balance between histone acetylases (HATs) and deacetylases (HDACs) influences gene expression patterns (13, 14).

Histone H3 a crucial component of the nucleosome, undergoes various post-translational modifications (PTMs) that impact gene regulation. The most widely studied PTM of H3 include acetylation of various lysine residues (K4, K9, K14, K18, K27, and K56) in the H3 tail. Acetylation of K9 and K27 on H3 facilitates transcription factor binding and gene activation. In contrast tri- methylation of the same K9 and K27 of H3 act as repressive signals associated with gene silencing (15). The balance between acetylation and methylation at specific lysine residues plays a critical role in regulating gene expression.

H3 phosphorylation is another dynamic PTM that plays a pivotal role in chromatin remodeling and gene expression (16, 17). Unlike acetylation and methylation, phosphorylation establishes interactions with other histone modifications and effector proteins serving as a platform for coordinating various regulatory events(18). Phosphorylation of H3 at serine 10 (H3S10) and serine 28 (H3S28) at gene promoters influence methylation and acetylation of neighboring lysines of H3 (19–22). The interplay between phosphorylation, acetylation and methylation create a dynamic regulatory work that fine-tune gene expression (23).

We recently demonstrated that IH increased acetylation and decreased methylation of H3 (24). Increased KDM6 (lysine demethylase) enzyme activity regulates H3K27 methylation status under the setting of IH (25). However, the mechanisms underlying IH-induced H3 acetylation have not been investigated. ROS (reactive oxygen species) dependent decrease in HDAC activity lead to increased HIF-1α acetylation by p300/CBP (CREB-binding protein) (HAT) in response to IH (26, 27). Based on these findings, we investigated the upstream regulators and the specific lysine residues that contribute to IH-mediated H3 hyper acetylation.

## METHODS

### Exposure of cell cultures to IH

Pheochromocytoma (PC) 12 cells (original clone from Dr. Green, Columbia University, New York, New York) (28) were cultured in Dulbecco’s modified Eagle’s medium (DMEM) supplemented with 5% fetal bovine serum (FBS) and 10% horse serum under 10% CO2 and 90% room air (20% O2) at 37°C (29). SH-SY5Y (human neuroblastoma; CRL-2266), MEF (mouse embryonic fibroblasts; SCRC-1008) and HEK293T (human embryonic kidney; CRL-3216) were commercially obtained from ATCC. Experiments were performed on cells starved overnight in serum-free DMEM medium. Cell cultures were exposed to IH (1.5% O2 for 30 sec followed by 20% O2 for 5 min at 37°C) as described (30). Ambient O2 levels in the IH chamber were monitored using an O2 analyzer (Alpha Omega Instruments). In experiments involving treatment with drugs, cells were treated with either drug or vehicle for 30 min before and during IH exposure.

### Transient transfections

For gain of function studies, PC12 cells were transiently transfected with overexpression plasmids pHDAC5, pHDAC3 and pNIPP1-GFP (addgene) or with control vector plasmids pcDNA3.1, pCMV or pGFP using TransIT-2020 (Mirus) transfection reagent. Briefly, cells were plated in 60mm tissue culture plates at a density of 2x10^6^ cells/plate in serum containing growth medium. DNA-TransIT reagent in the ratio of 1:3 was added after 24hrs, Serum starved cells were exposed to IH after 36hrs of transfection.

For loss of function studies, recombinant lentivirus was generated by transfection of HEK293T cells with the third-generation packaging plasmids, *pMDLg*/*pRRE*, pRSV-Rev, pMD2-G (#12251, #12253,#12259 respectively from Addgene) and piLenti-siRNA-GFP encoding for p300 (#C044- G), or CBP (#168160960395) as well as non-targeting control siRNA (# LV015-G) from Applied Biological Materials (abm; Richmond, BC, Canada), using Trans IT-Virus GEN transfection Reagent (Mirus). Lentiviral supernatants were harvested, passed through a 0.45-μm filter, and aliquots were frozen at -80°C. PC12 cells transfected with lentivirus were cultured in complete medium for 48 h before exposure to IH. Transfection efficiency as determined by counting number of green fluorescent positive cells treated with GFP lentiviral vector was between 80-90%.

### Immunoblot assays

Cell extracts from PC12 cells were fractionated by polyacrylamide-SDS gel electrophoresis. Immunoblots were probed with the primary antibodies (see table 1) followed by corresponding HRP-conjugated secondary antibody and detected using Clarity Western ECL substrate kit (Bio-Rad, Hercules, CA). Immunoblots were quantified using Image studio software by Odyssey Fc imaging system (LI-COR, Lincoln, NE). Raw pixel data for each protein is normalized to loading control (tubulin) and expressed as fold change relative to vehicle treated controls under normoxia.

**TABLE 1:**
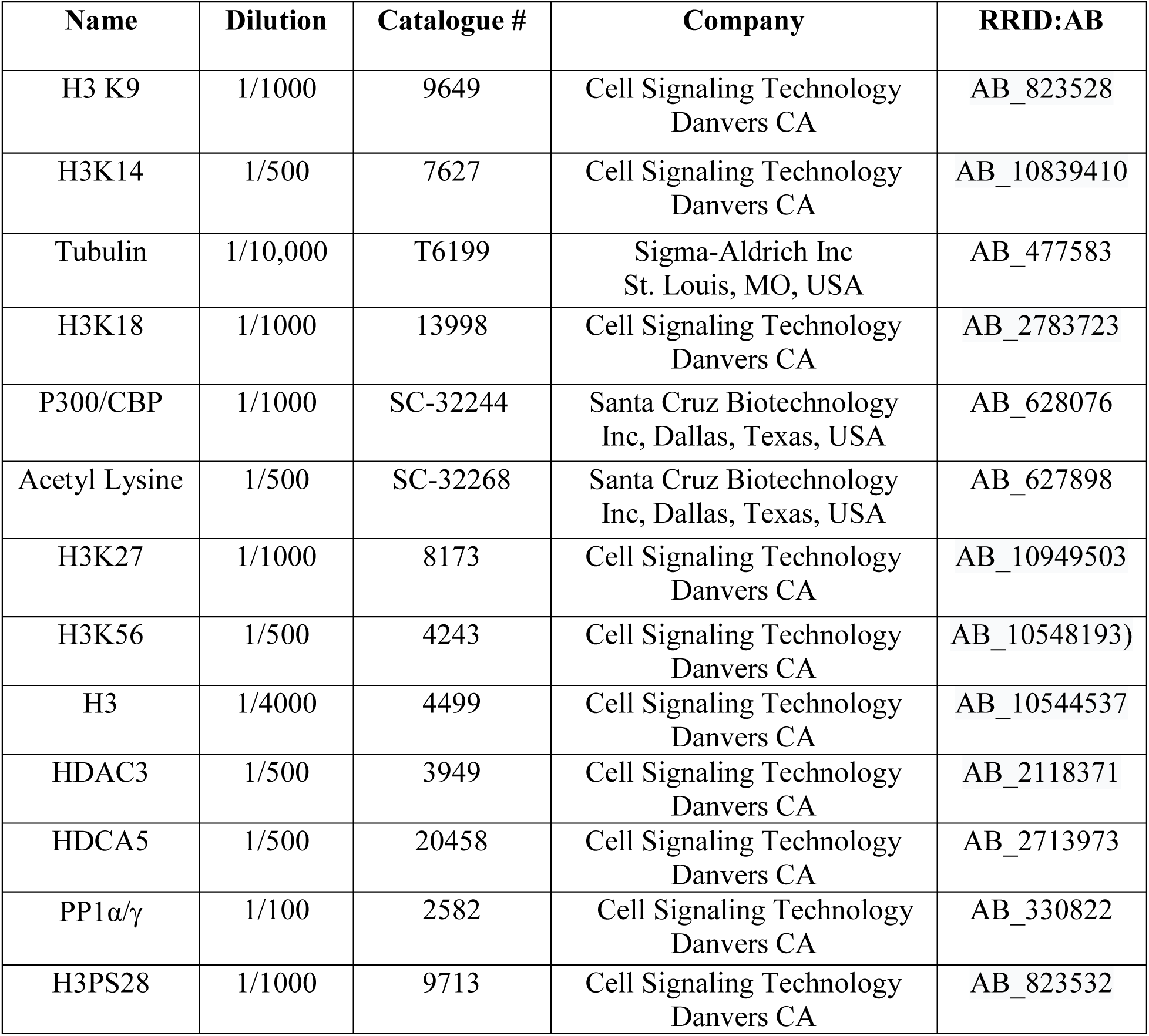
Primary antibodies information used for the study.

**Statistical analysis:** Data are expressed as mean ± S.E.M from 3-6 independent cell culture experiments, per group. Inter-group differences were assessed by one-way ANOVA with post hoc analysis using Tukey’s test, as appropriate. Values of *p* < 0.05 were considered statistically significant.

## RESULTS

### IH increased H3 acetylation in different cell types

Pheochromocytoma (PC) 12 cells cultures serve as a model system for O2-sensitive catecholamine-producing cells of adrenal medulla (AM) (31). Exposure of PC12 cell cultures to 60 cycles of IH (alternate cycles of hypoxia for 30 sec and reoxygenation for 5mts) increased global H3 acetylation (24). To assess specificity, H3 acetylation was analyzed in other cell lines like SH-SY5Y (human neuroblastoma), MEF (mouse embryonic fibroblasts) and HEK293T (human embryonic kidney cells) exposed to 60 cycles of IH. As shown in **Fig 1a**, IH increased H3 acetylation in all cell lines studied albeit to varying degrees. We next analyzed the time course of IH-induced H3 acetylation. PC12 cells exposed to increasing cycles of IH exhibited progressive H3 acetylation with a significant increase occurring after 30 cycles (**Fig. 1B**). Remarkably, IH-induced H3 acetylation was reversible. Within 16h after recovery in room air, the elevated H3 acetylation levels returned to base line (**Fig. 1B**).

**Figure 1:**
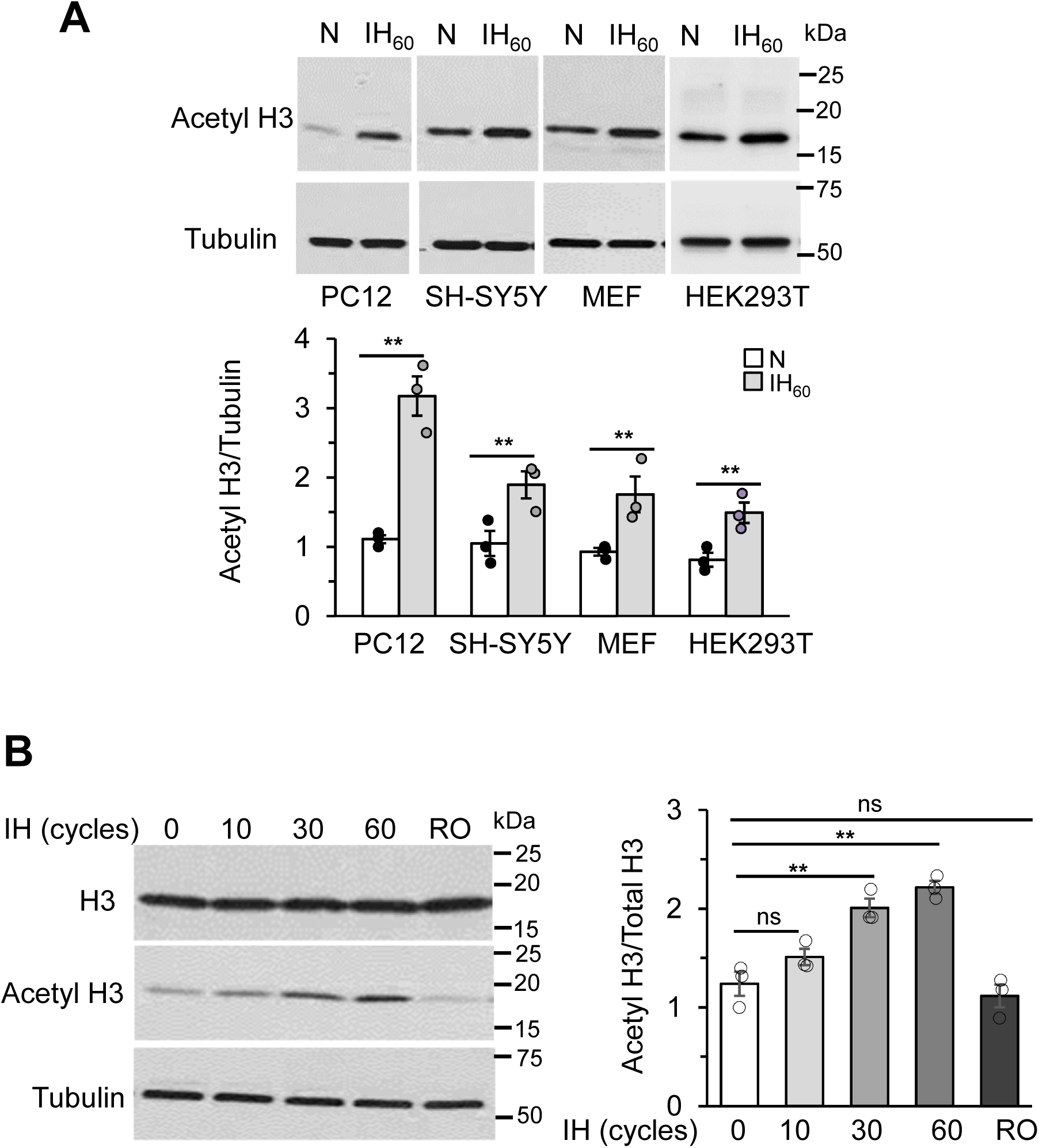
H3 acetylation in different cell lines and time course of H3 acetylation in response to IH. **A)**. *Top Panel*: Representative example of an immunoblot showing H3 acetylation in lysates from room air and IH-exposed PC12, SH-SY5Y, MEF and HEK293T cells. *Bottom panel*: Quantitative analysis of the blots by image J **B)** *Left panel*: H3 acetylation, total H3 protein and tubulin expression in PC12 cells exposed to normoxia and increasing cycles of IH. *Right Panel*: Quantitative analysis of the blots by image J. Data was normalized to tubulin loading control and expressed as fold change relative to control (normoixa) (mean ± SEM; n=3-4). \*\**p*≤0.05; ns = not significant (*p*>0.05) as determined by One-way ANOVA with post-hoc Turkey HSD test.

### XO derived ROS contributes to IH-increased H3 acetylation

ROS signaling mediates cellular and systemic responses to IH (32). Given that IH increases ROS levels and ROS are potent post translational regulators of proteins (27), we hypothesized that ROS mediates IH induced increase in H3 acetylation. To assess this possibility, PC12 cells exposed to IH, were treated with MnTmPyP (Mn(III)tetrakis(1-methyl-4-pyridyl) porphyrin), a membrane permeable anti-oxidant. IH-induced increase in H3 acetylation was inhibited in the presence of 50μM MnTmPyP (**Figure 2A**). Conversely, treating control room air cultured cells with Xanthine (250ug) and increasing concentrations of Xanthine oxide (XO, 0.01-0.1U/ml), to generate ROS or with tert-butyl hydroperoxide (TBH,10-100uM), an established ROS mimetic, increased H3 acetylation dose-dependently (**Fig 2B-C**). We next determined the source of ROS which mediate the effects of IH on H3 acetylation. Our earlier studies identified xanthine oxidase (XO) and NADPH oxidase (NOX) as major ROS sources during IH (33–36). To determine which of these two oxidases contribute to H3 acetylation by IH, cells were treated with 50µM allopurinol, an established inhibitor of XO, or 20µM VAS2870, a pan inhibitor of Nox isoforms, and exposed to IH or room air. Allopurinol blocked IH-induced H3 acetylation while VAS2870 had no effect (**Fig 2D**). These results highlight the role of XO generated ROS in regulating H3 acetylation during IH.

**Figure 2:**
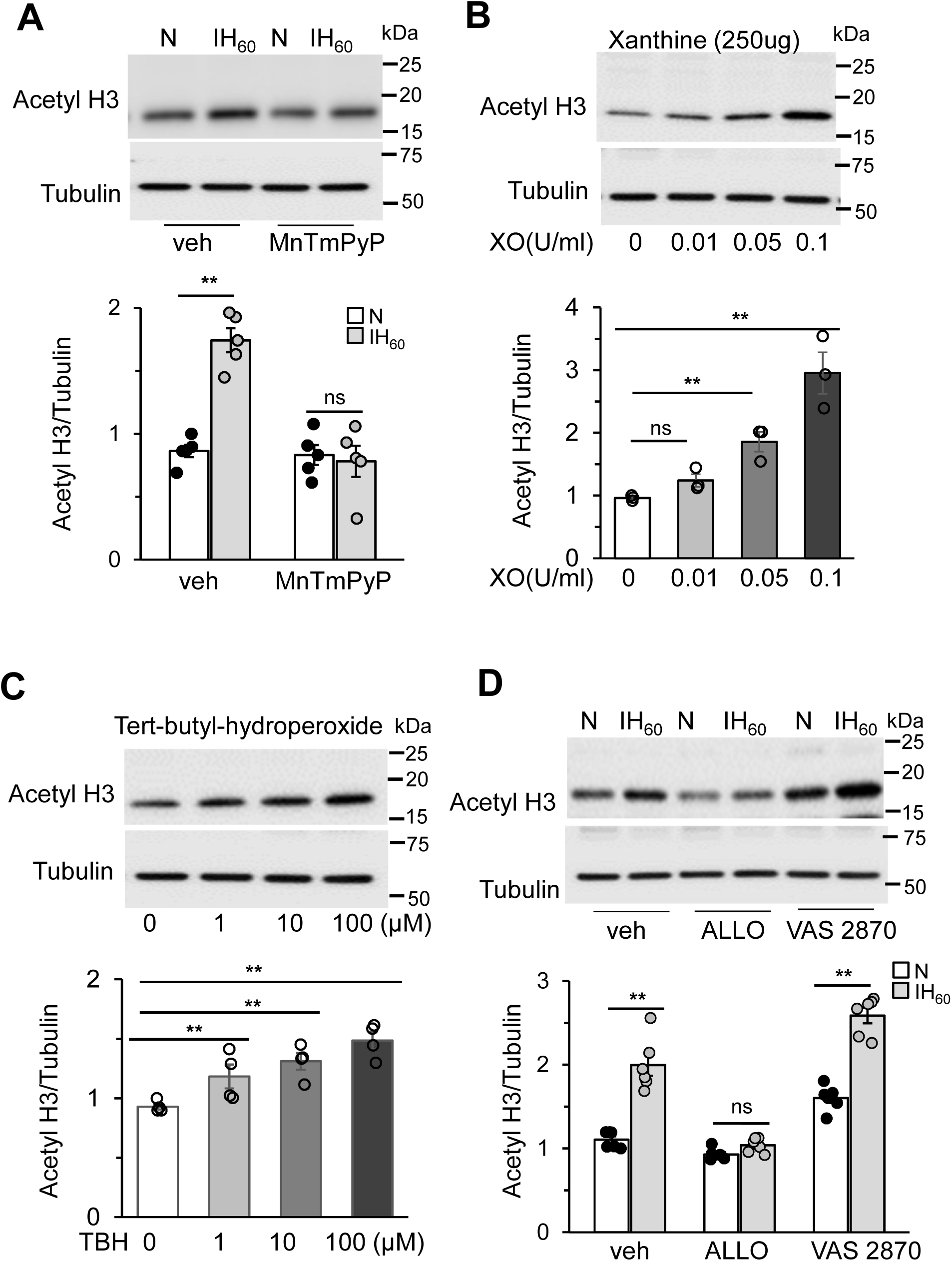
ROS generated by Xanthine oxidase (XO) regulate H3 hyperacetylation in response to IH. **A**) *Top panel*: Acetylated H3 protein expression in room air (N) and IH60 exposed cells treated with vehicle or MnTmPyP (50µM). *Bottom panel*: Quantitative analysis of the blots using Image studio by Odyssey Fc imaging system (n=4). **B –C)** Representative immunoblots (*top panel*) and quantitative analysis of the blots (*bottom panel*) of acetylated H3 protein expression in PC12 cells treated with different concentrations of ROS mimetics (Xanthine/Xanthine oxidase (XO)) and Tertiary butyl hydroperoxide (TBH) (mean ± SEM; n=4) **D)** *Top panel* Representative immunoblot showing acetylated H3 protein expression and tubulin as loading control in cell lysates from room air (N) and IH60 exposed PC12 cells treated with Allopurinol (XO inhibitor; 50µM) or VAS2870 (NOX inhibitor; 20µM). *Bottom panel*: Quantitative analysis of the blots (mean ± SEM; n=3) using Image Studio by Odyssey Fc imaging system. H3 acetylated protein expression was normalized to tubulin and expressed relative to vehicle treated control cells exposed to room air (N). \*\**p*≤0.05; ns = not significant (*p*>0.05) as determined by Mann-Whitney test.

### Reduced HDAC3 contribute to IH-induced increase in H3 acetylation

We then sought to determine the mechanism by which ROS increases H3 acetylation in response to IH. Acetylation status of H3 is dictated by a balance between HATs and HDACs. IH increased ROS generation contributes to decreased HDAC activity (24). We therefore assessed whether reduced HDAC activity is sufficient to increase H3 acetylation. This possibility was tested by determining H3 acetylation in PC12 cells treated with 50nM Trichostatin A (TSA), a pan inhibitor of HDACs. TSA treatment significantly increased H3 acetylation under normoxic conditions and IH had no further effect (**Fig 3A**). Given that IH reduced HDAC activity was due to proteasomal degradation of HDAC5 and HDAC3 proteins (24), we examined the relative contribution of HDAC3 and HDAC5 to IH-induced H3 acetylation. To this end, cells were transfected with over expression plasmids encoding either HDAC3 or HDAC5, and then exposed to IH60 or normoxia (control). Over expression of HDAC3 blocked the reduced HDAC3 expression as well as H3 acetylation characteristic to IH (**Fig. 3B**). Overexpression of HDAC5 had no effect on IH-induced H3 acetylation but blocked IH-induced decrease in HDAC5 expression but (**Fig. 3C**). These findings demonstrate that reduced HDAC3 but not HDAC5 contribute to increased H3 acetylation by IH.

**Figure 3:**
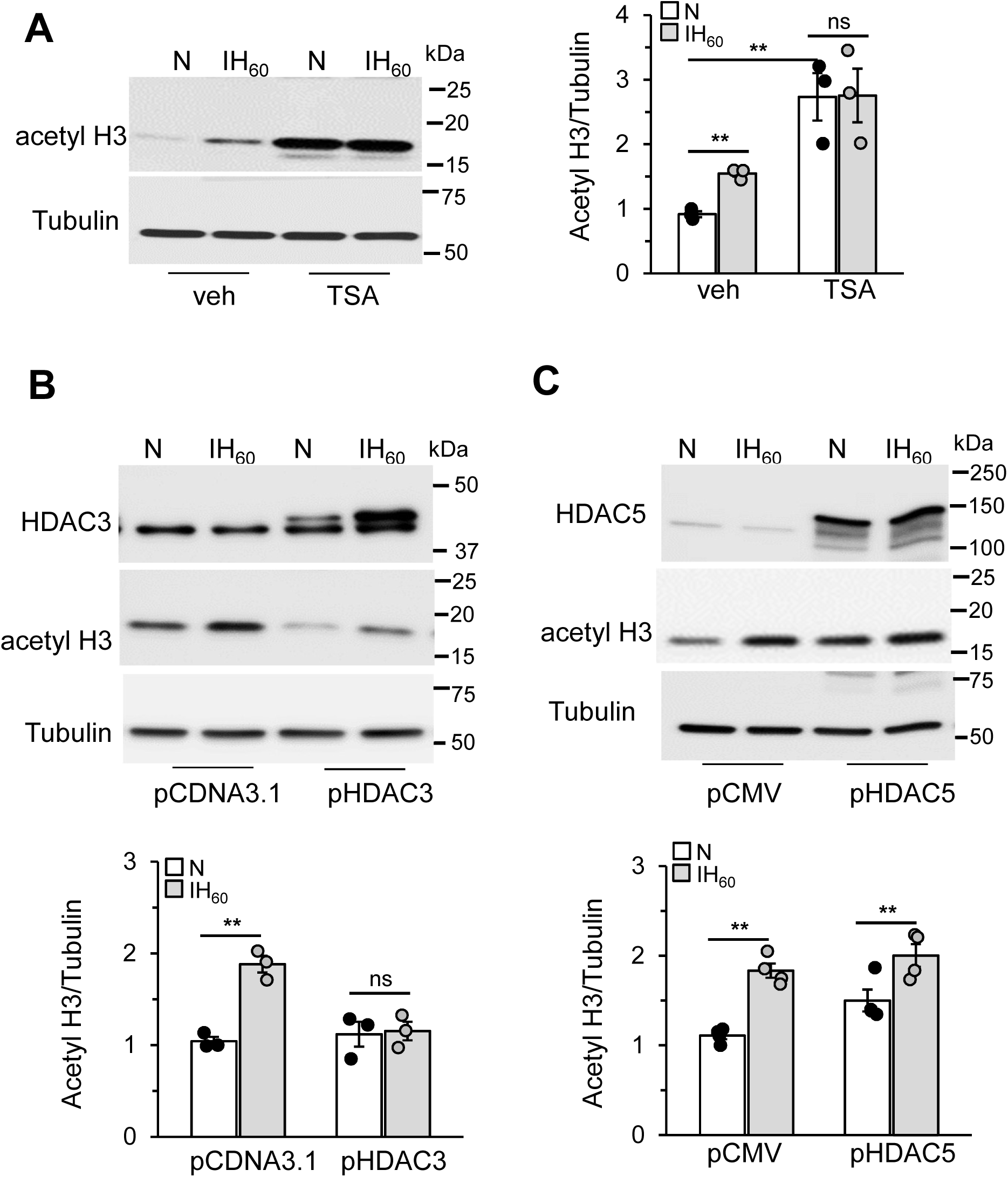
Role of HDACs in IH-induced H3 hyperacetylation. **A**): *Left panel* Representative immunoblot showing H3 acetylation with tubulin as a loading control in cell lysates from room air (N) and IH-exposed cells treated with vehicle or TSA (50nM for 7hrs). *Right Panel*: Quantitative analysis of the blots. **B &C)** *Top Panel*: H3 acetylation, HDAC3, and HDAC5 protein expression with tubulin as loading control in cell lysates from room air (N) and IH-exposed PC12 cells transfected with overexpression plasmids encoding either HDAC3, HDAC5, or control vector pcDNA. *Bottom panel*: Quantitative analysis of the blots by Image Studio software from Odyssey Fc imaging system (LI-COR, Lincoln, NE) (mean ± SEM; n=3-4). \*\**p*≤0.05; ns = not significant (*p*>0.05) as determined by One-way ANOVA with post- hoc Turkey HSD test.

### H3 acetylation by IH requires p300 protein

IH increased HAT proteins, p300/CBP (TIP 60 family) but not PCAF and GCN5 proteins (GNAT family) (26). To assess if p300/CBP contributes to H3 acetylation by IH, cells exposed to IH or room air were treated with CTK7A, a water-soluble, cell permeable inhibitor of p300/CBP (37) or CPTH6 (38), inhibitor of GCN5 and pCAF. 25µM CPTH6 had no significant effect on p300/CBP or H3 acetylation proteins. On the other hand, 25µM CTK7A inhibited both p300/CBP protein level and H3 acetylation (**Fig 4A**) in response to IH. To ascertain the individual role of p300 and CBP in IH induced H3 acetylation, cells were treated with scrambled lentiviral siRNA or siRNA directed to p300 or CBP. Compared to cells treated with scrambled siRNA, p300 siRNA treated cells exposed to IH showed reduced p300/CBP protein level and H3 acetylation, whereas CBP siRNA-treated cells showed an absence of increased p300/CBP protein, but elevated H3 acetylated protein (**Fig. 4B**). These results indicate that the increased H3 acetylation in response to IH is mediated by p300 but not CBP.

**Figure 4:**
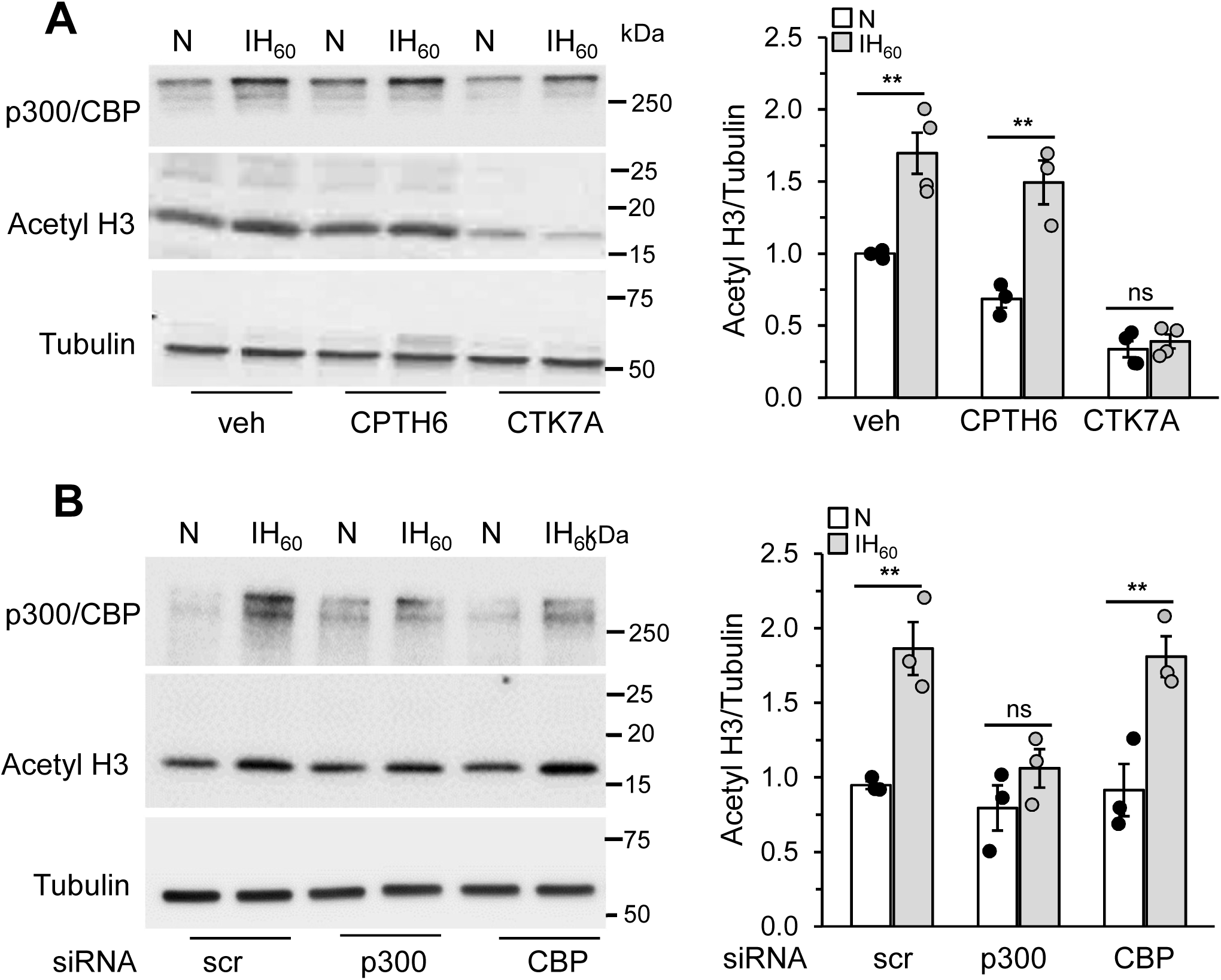
p300 contributes to IH-augmented H3 acetylation. **A):** *Left panel* Representative immunoblot showing p300/CBP and acetylated H3 protein expression with tubulin as loading control in PC12 cells treated with vehicle or 50 µM of CPTH6 (inhibitor of pCAF and GCN5) or 25µM of CTK7A (inhibitor of p300/CBP) and exposed to IH60 or normoxia (N). *Right Panel:* Quantitative analysis of the immunoblots. **B)** p300/CBP and acetylated H3 protein expression with tubulin as loading control in PC12 cells transfected with scrambled or p300 or CBP lentiviral siRNA and exposed to IH60 or normoxia (N). *Right Panel:* Quantitative analysis of the immunoblots as analyzed by Image Studio software from Odyssey Fc imaging system. Raw pixel values were normalized to tubulin loading control and expressed as fold change relative to normoxia controls, (mean ± SEM; n=3-4). \*\**p*≤0.05; ns = not significant (*p*>0.05) as determined by One-way ANOVA with post-hoc Turkey HSD test.

### Role of phosphatases in increased H3 acetylation by IH

H3 acetylation, in addition to being regulated by HDACs and HATs, is also influenced by H3 phosphorylation, which plays a crucial role in chromatin remodeling and gene expression (16).

H3 phosphorylation is regulated by dynamic equilibrium between nuclear protein kinases including Aurora B kinase and mitogen or stress activated kinases (MSK1 and MSK2) which phosphorylate H3 and nuclear protein phosphatases (PP1 and PP2) which catalyze dephosphorylation of H3 (39, 40). If increased phosphorylation of H3 is necessary to induce acetylation, then inhibition of Aurora B and MSK kinases should block IH induced H3 acetylation. However treating cells with Barasertib (Aurora B kinase inhibitors) (41), SB202190 (p38 inhibitor) (42) or SB747651A (MSK1 inhibitor)(43) did not block IH-induced H3 acetylation. (**Suppl Fig 1A**). To assess whether the increase in H3 acetylation involve phosphatases, IH exposed cells were treated with a phosphatase inhibitor cocktail (PIC) containing three inhibitors including: Calyculin A, an inhibitor of protein phosphatases 1 (PP1) and 2A (PP2A); Cantharidin an inhibitor of PP2A; and (-)-p-Bromolevamisole oxalate (BTO), an inhibitor of L-isoforms of alkaline phosphatases. PIC blocked IH-induced H3 acetylation without affecting total H3 protein levels (**Fig 5A**). To identify the specific phosphatase(s) contributing to H3 acetylation, cells cultured in room air were treated with different concentrations of either BTO, Cantharidin or Calyculin A, and H3 acetylation were subsequently analyzed. Calyculin A decreased the expression of acetylated H3 protein, whereas neither BTO nor cantharidin had any effect. (**Suppl Fig 1B**).

**Figure 5:**
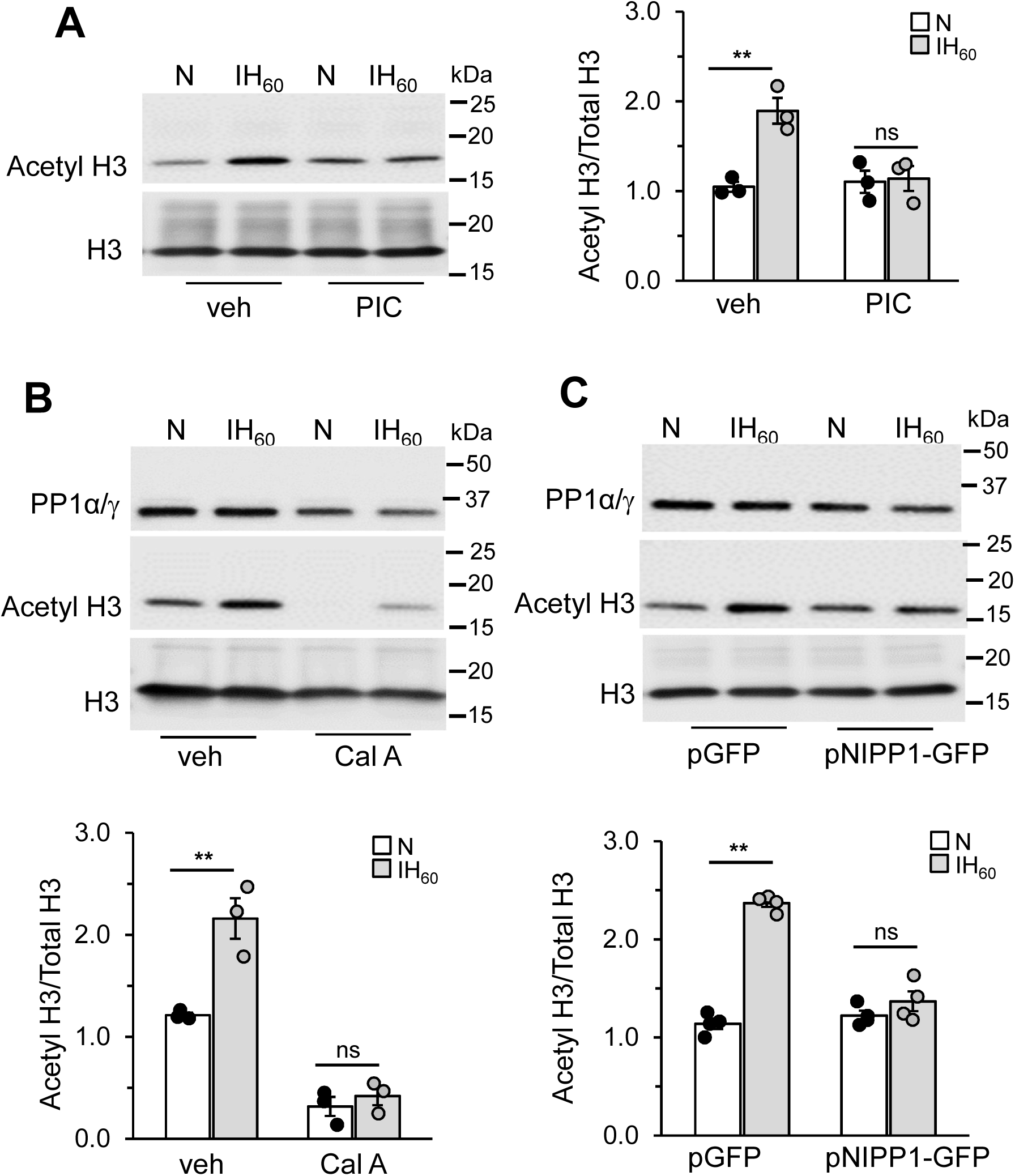
Protein phosphatase 1 (PP1α/**γ**) dependent dephosphorylation results in H3 acetylation by IH. **A**) *Left panel*: Acetylated and total H3 protein in lysates from room air (N) and IH60-exposed cells treated with vehicle or phosphatase cocktail inhibitor (PIC) (1/100 dilution) or Calyculin A (10nM) (n=3). *Right Panel:* Quantitative analysis of the blots using Image Studio. **B)** *Top pane:l* Acetylated H3, total H3 and PP1α/γ protein in lysates from room air (N) and IH60-exposed cells treated with vehicle or Calyculin A (10nM). *Bottom panel:* Quantitative analysis of the blots (n=3). **C)** *Top panel:* Representative immunoblot showing acetylated H3, PP1α/γ protein and tubulin as loading control in lysates from room air (N) and IH60exposed cells transfected with either NIPPI- GFP or control vector GFP plasmids *Bottom panel:* Quantitative analysis of the blots (n=3-4). \*\**p*≤0.05; ns = not significant (*p*>0.05) as determined by one-way ANOVA with post-hoc Turkey HSD test.

Calyculin A binds and inhibits both PP1 and PP2A catalytic subunits with similar IC50 values (44). Previous studies showed that Calyculin A reduced PP1α/γ protein abundance without affecting PP2A-A, PP2A-B and PP2A-C proteins in both control (room air treated) and IH treated cells (27). Calyculin A treatment decreased PP1α/γ protein expression as reported previously and blocked IH-induced increase in acetylated H3 without any effect on total H3 protein (**Fig 5B**) suggesting a role for PP1α/γ in H3 acetylation by IH. <u>N</u>uclear inhibitor of <u>PP1</u> (NIPP1) is an endogenous PP1 inhibitor (45) and PP1α/γ is activated by disassociating from NIPP1 complex . If PP1α/γ contributes to H3 hyperacetylation by IH, then inhibiting PP1 activity with ectopic expression of NIPP1 should block the H3 acetylation in IH treated cells. To assess this possibility, cells were transfected with a GFP tagged NIPP1 plasmid (46) and exposed to IH. As shown in **Fig 5C**, ectopic expression of GFP tagged NIPPI reduced PP1α/γ protein levels and prevented IH-induced H3 acetylation in both control and IH exposed cells. Taken together these findings demonstrate that inhibition of PP1 activity pharmacologically (Calyculin A) or with ectopic expression of NIPP1 blocks H3 acetylation increase by IH, suggesting a role for phosphorylation.

### H3 S28 phosphorylation influences H3 K27 acetylation in response to IH

The above results prompted us to analyze if the effects of phosphatase inhibitors are due to direct alteration of phosphorylation status of H3 by IH. H3 can be phosphorylated at H3T6, S10, T11 and S28 adjacent to K9, K27, and K14 residues (16). We therefore first examined which lysine residues account for acetylation of H3 by IH. To this end, acetylation of K56, K27, K18, K14 and K9 were determined in cell extracts of IH-treated cells by immunoblot assay. IH led to increased acetylation of K27, K18 and K14 on H3. However, acetylation levels of K56, and K9 were not altered by IH (**Fig 6A**). Treatment of cells with TSA (pan HDAC inhibitor), under normoxic conditions (room air treated), robustly increased acetylation of all lysine residues (**Fig 6A**).

**Figure 6:**
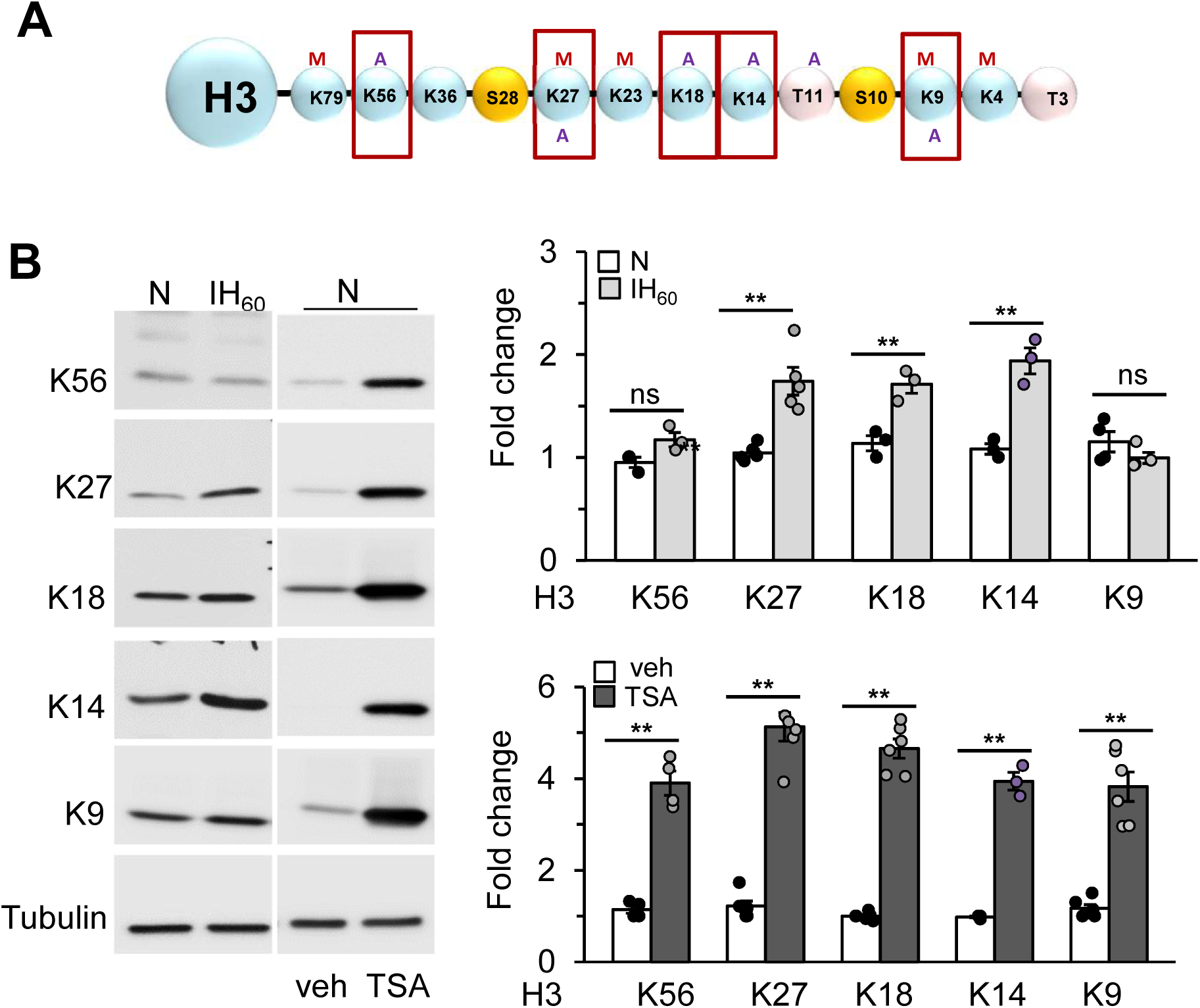
H3 lysines acetylated in response to IH. **(A)** Schematic diagram showing the acetylation (A) and methylation (M) of lysines of H3 N-terminal tail. **B)** *Left panel*: Acetylation of H3K9, H3K14, H3K18, H3K27 and H3K56 in lysates of PC12 cells from room air (N), IH-exposed as well as veh or TSA (50nM for 7hrs) treated PC12 cells. *Right panel:* Quantitative analysis of the blots (n=3-4). Values were normalized to tubulin loading control and expressed as fold change to the corresponding room air exposed cells (N)

We chose to focus on K27 acetylation to analyze the crosstalk between acetylation and phosphorylation for the following reasons: 1) K27 can be both acetylated and methylated, 2) IH increased K27 acetylation, and decreased K27 methylation (25), and 4) phosphorylation of H3 at S28 influences acetylation of K27 (19, 21).

Since phosphatase inhibitors PIC and Calyculin inhibited global H3 acetylation, we analyzed the effect of these drugs on S28 phosphorylation and H3K27 acetylation in PC12ells exposed to normoxia or IH. IH alone had no significant effect on S28 phosphorylation but both Calyculin and PIC increased S28 phosphorylation under normoxic conditions compared to vehicle treated cells (**Fig 7A**). Calyculin treatment had no further effect in response to IH. In contrast PIC increased S28 phosphorylation in cell lysates exposed to IH compared to normoxia (**Fig 7A**). Increased S28 phosphorylation with Calcyculin and PIC treatment correlated with decreased K27 acetylation (**Fig 7A**) with no change in total H3 protein. To further confirm the correlation between S28 phosphorylation and K27 acetylation, PC12 cells were exposed to IH or room air (controls), with and without CTK7A treatment which inhibited global H3acetylation. CTK7A treatment increased S28 phosphorylation and decreased H3K27 acetylation in cell lysates of both room air and IH exposed cells compared to vehicle treated cells (**Fig 7B**).

**Figure 7:**
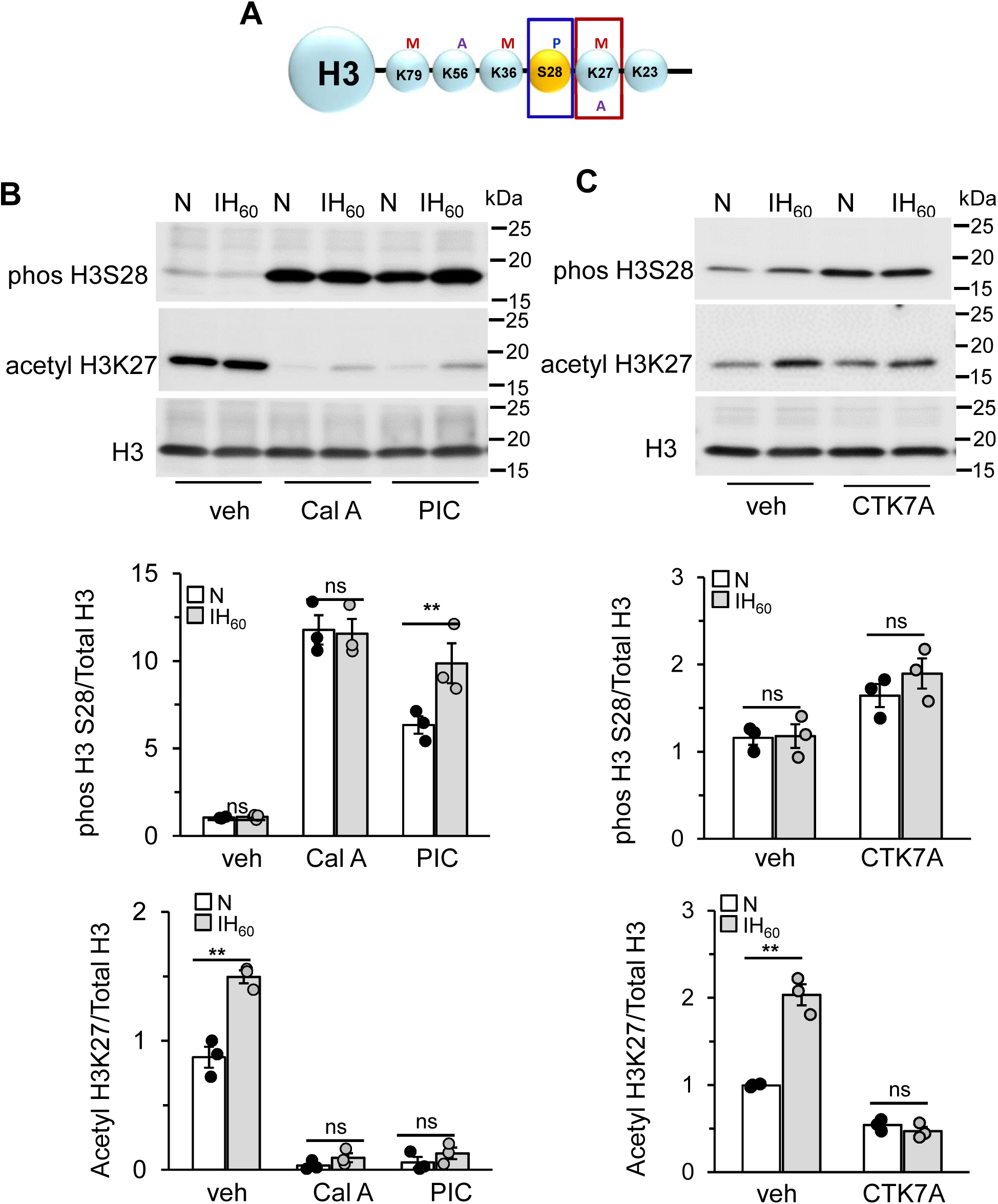
H3 S28 phosphorylation and H3 K27 acetylation in response to IH. **A)** Schematic diagram showing the relative position of H3S28 phosphorylation (P) to H3K27 acetylation (A) and H3K27 methylation (M). **B & C)** *Top panels*: H3S28 phosphorylation, acetylated H3K27 and total H3 protein expression in lysates from room air (N) and IH60-exposed PC12 cells treated with **B)** vehicle or Calyculin A(10nM) or PIC (1/100 dilution) or **C)** vehicle or CTK7A. *Bottom panels*: Quantitative analysis of the blots expressed as ratio of phosphorylated H3S28 and acetylated H327 to total H3 (n =3).

## DISCUSSION

Major findings of the current study are 1) IH evoked HDAC inhibition resulted in increased global H3 acetylation by p300/CBP protein (HAT) without affecting total H3 protein expression, 2) IH- induced H3 acetylation is mediated by XO generated ROS, 3) PP1α/γ, a protein phosphate, inhibitor blocked IH-induced H3 acetylation, 4) Acetylation of K14, K18 and K27 contributed to overall increase in H3 acetylation induced by IH, and 5) IH increased H3K27 acetylation, was regulated by phosphorylation of adjacent serine 28 residue (H3S28). Our study demonstrates a novel crosstalk between H3 acetylation and phosphorylation post-translational modifications, highlighting the complexity of H3 function in modulating the epigenome during IH.

ROS are the primary mediators of systemic and cellular responses to IH in OSA patients and in experimental rodent models treated with IH (47–49). ROS scavenger prevented H3 acetylation by IH and ROS mimetics, increased global H3 acetylation (similar to IH) in control cells exposed to room air demonstrating that ROS mediate IH-induced H3 acetylation.

Pharmacological blockade of Xanthine oxidase (XO) inhibited H3 acetylation by IH implicating XO as the major source of ROS mediating IH’s effects on H3 acetylation.

How might ROS contribute to the increased H3 acetylation by IH? ROS dependent decrease in HDAC enzyme activity was associated with increased HIF-1α acetylation in response to IH(24). Consistent with these findings, decreased HDAC activity was also associated with increased H3 acetylation. Thus IH-induced ROS likely contribute to H3 acetylation by regulating HDAC activity. Alternatively H3, the only histone, which contains cysteine’s, has been shown to sense redox changes through S-glutathionylation (50), which leads to a more open chromatin structure. We have previously shown that IH induced perturbation of redox homeostasis involved glutathionylation of complex 1 subunit proteins (51). Whether IH alters glutathionylation of H3 through ROS signaling remains an open question.

IH reduced HDAC activity was associated with decreased protein abundance of HDAC5 (class II) and HDAC3 (class I) proteins (24). Overexpression of HDAC3 alone was sufficient to block IH-induced H3 acetylation, while overexpression of HDAC5 did not alter H3 acetylation by IH. This is in contrast to IH-induced HIF-1α acetylation which is regulated by HDAC5 but not HDAC3 (24). These results suggest that individual HDACs play distinct roles in the context of IH and highlight the complexity of HDAC-mediated epigenetic regulation during IH, emphasizing the importance of specific HDAC isoforms.

Both HIF-1 and H3 are acetylated in response to IH, however there are distinct differences which are as follows: 1) acetylation stabilizes HIF-1α but has no effect on H3 protein, 2) HDAC3 inhibition is involved in H3 acetylation while HDAC5 is involved in acetylation of HIF-1α and 3) XO generated ROS contributes to H3 acetylation while NADPH oxidases regulate HIF-1 acetylation. In spite of the differences observed above, our results demonstrated that similar to HIF-1α, H3 is acetylated by p300 the most widely studied member of the HAT family. Based on the differences and our previous finding that XO generated ROS is the initial trigger for NADPH oxidase activation (33), we speculate that H3 acetylation precedes HIF-1α acetylation. It is conceivable that H3 acetylation by p300 may not only play a role in facilitating HIF-1α acetylation but also HIF-1α accessibility to target gene promoters by opening chromatin promoter region of target genes, a possibility that requires additional studies.

H3 phosphorylation, a dynamic post-translational modification (PTM) establishes interactions with effector proteins, and influences the methylation and acetylation of neighboring lysines on H3 (16, 18). Interestingly, inhibitors of nuclear kinases, Aurora B and MSK1which phosphorylate H3 were ineffective in blocking IH-induced H3 acetylation. However, phosphatases (PP1 and PP2) inhibitors or overexpression of endogenous PP1 inhibitor (NIPP1) blocked H3 acetylation by IH, implicating phosphatase PP1α/γ as a regulator of IH-induced H3 acetylation. We have previously shown that PP1α/γ regulates ROS-dependent degradation of HDAC5 by IH (27). These findings suggest that phosphatases can influence H3 acetylation via two distinct mechanisms, indirectly by regulating effector proteins like HDACs or directly by modulating H3 phosphorylation status.

Activation of specific target genes often involves coordinated modification of H3 including phosphorylation and acetylation on adjacent residues within the H3 tail (21, 23). H3 can be acetylated at multiple lysine positions including Lys 4, 9, 14, 18, 27, and 56. Our studies showed that H3K27 but not H3K9 is one of the lysines acetylated by IH. Juxtaposed to H3K27 is the widely conserved H3S28 phosphorylation site shown to influence H3K27 acetylation. We could not detect significant differences in H3S28 phosphorylation in response to IH. This could be due to sensitivity of the antibody as studies have shown inducible histone H3 phosphorylation is rapid, transient and is a very small fraction of total H3. Interestingly, phosphatase inhibitor treatments increased S28 phosphorylation significantly which correlated with decreased K27 acetylation suggesting a regulatory relationship between these modifications. The apparent loss of H3K27 acetylation with PIC treatment could be due to epitope masking or occlusion. When the adjacent S28 residue is heavily phosphorylated the H3K27 acetylated antibody may not recognize its epitope effectively. However, treatment of cells with CTK7A which inhibited K27 acetylation increased S28 phosphorylation eliminating the potential epitope occlusion problem and providing a link between H3 phosphorylation and acetylation. Clearly additional studies are needed to explore the relationship between these modifications at gene promoters using ChiP assays to see if these two modifications are two independent but spatially linked events or one modification is a prerequisite for the other. Nonetheless, our results suggests an unacknowledged function of unphoshporylated H3S28 in H3K27 acetylation. Although H3K9 acetylation remained unaltered with IH, whether other serine phosphorylation sites including H3S10 adjacent to K9 also contribute to H3K27 acetylation by IH remains to be investigated.

We have previously shown that IH decreases methylation of H3K27 due to increased KDM6 (demethylase) enzyme activity (25), suggesting IH induces a switch from methylation to acetylation at H3K27 for transcriptional activation. Phosphorylation of H3 is required to enable proper methylation of H3 to establish gene silencing as well as to promote removal of the repressive methyl marks on H3 by demethylases (16). It is therefore likely, H3S28 phosphorylation is required for H3K27 methylation for repression of genes under normoxia. In response to IH H3S28phophsorylation could promote removal of K27methylation marks, facilitating K27acetylation a possibility that requires additional studies and is beyond the scope of the current investigation. Our findings provide valuable insights into the complex interplay between H3S28 phosphorylation, H3K27 acetylation and H3K27methylation in modulating gene expression and as potential therapeutic targets in response to IH.

## Authors Contribution

J.N. conceived and designed the experiments. N. W, and M.H performed experiments and analyzed the data. J.N wrote and N.R.P edited the manuscript.

## Acknowledgement

This work was supported by the National Institutes of Health grant P01-HL144454.

