## supplemental fig 1 and 2 for "Reactive Oxygen species dependent increase in H3K27 acetylation by intermittent hypoxia is regulated by H3S28 phosphorylation"

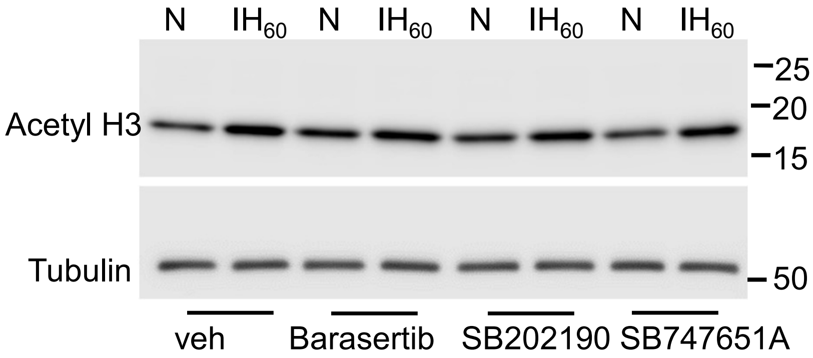


Fig S1: Role of protein kinases in IH induced H3 acetylation. Representative immunoblot showing acetylated H3 and tubulin protein expression in PC12 cells exposed to room air (N) or $IH_{60}$ treated with inhibitors of Aurora B kinase (Barasertib, 10μM), p38 kinase (SB202190, 10μM) or MSK1 (SB747651A, 10μM).

Supplementary Fig S1

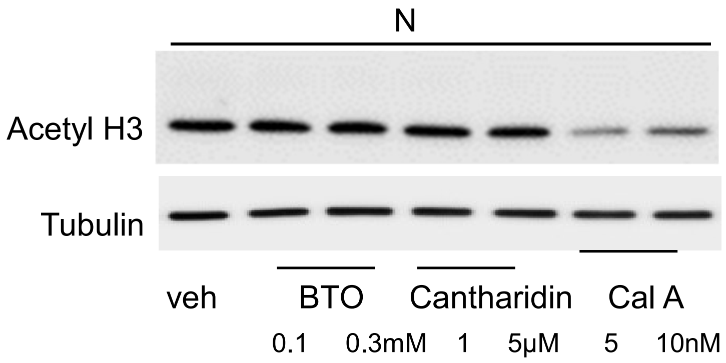


Fig S2: Effect of phosphatase inhibitors on IH-induced H3 acetylation. Acetylated H3 and tubulin protein expression in PC12 cells treated with two different concentrations of p-Bromolevamisole oxalate (BTO), Cantharidin and Calyculin A (Cal A), individual components of PIC in room air (n=3).

Supplementary Fig S2
